# Acetylated α-tubulin residue K394 regulates microtubule stability to shape the growth of axon terminals

**DOI:** 10.1101/2021.04.01.438108

**Authors:** Harriet A. J. Saunders, Dena M. Johnson-Schlitz, Brian V. Jenkins, Peter J. Volkert, Sihui Z. Yang, Jill Wildonger

## Abstract

Microtubules are essential to neuron shape and function. Therefore, the stability of the microtubule cytoskeleton must be carefully regulated. Acetylation of tubulin has the potential to directly tune microtubule stability, and proteomic studies have identified several acetylation sites in α-tubulin. This includes the highly conserved residue lysine 394 (K394), which is located at the αβ-tubulin dimer interface. Using a fly model, we show that α-tubulin K394 is acetylated in the nervous system and is an essential residue. We found that an acetylation-blocking mutation in endogenous α-tubulin, K394R, perturbs the synaptic morphogenesis of motoneurons by reducing microtubule stability. Intriguingly, the K394R mutation has opposite effects on the growth of two functionally and morphologically distinct motoneurons, revealing neuron-type-specific responses when microtubule stability is altered. Eliminating the deacetylase HDAC6 increases K394 acetylation, and the over-expression of HDAC6 reduces microtubule stability similar to the K394 mutant. Thus, our findings implicate α-tubulin K394 and its acetylation in the regulation of microtubule stability and suggest that HDAC6 regulates K394 acetylation during synaptic morphogenesis.

## Introduction

The microtubule cytoskeleton is comprised of stable and dynamic microtubules that are integral to creating and maintaining neuronal morphology. Stable microtubules neither grow nor shrink, whereas dynamic microtubules undergo bouts of slow growth followed by rapid shrinkage (called “catastrophe”) as dimers of α- and β-tubulin are added or removed from microtubule ends. It is well-known that disrupting microtubules alters neuronal structure and function (Penazzi et al., 2016; Bodaleo and Gonzalez-Billault, 2016). Thus, microtubule stability and dynamics must be carefully regulated to attain proper neuronal morphology, although how this is achieved remains largely unknown.

In neurons and other cells, microtubule stability is regulated by a combination of factors, including the intrinsic properties of tubulin, proteins that interact with tubulin and microtubules, and post-translational modifications. Post-translational modifications, such as acetylation, have long been candidates to regulate microtubule stability and dynamics in cells. Although multiple sites of acetylation have been identified in tubulin, only a single non-essential site in α-tubulin, lysine 40 (K40), has been well characterised (Janke and Magiera, 2020). Acetylation of α-tubulin K40 correlates with stable microtubules in cells but does not itself regulate microtubule stability or dynamics. Rather, K40 acetylation enables microtubules, particularly long-lived microtubules, to withstand mechanical stresses, which may explain its correlation with stable microtubules (Portran et al., 2017; Xu et al., 2017). Although there is potential for microtubule stability to be controlled by acetylation of other sites in α-tubulin, this has not yet been explored.

Proteomic studies using tissue from mammals and flies have revealed that at least twelve conserved sites in α-tubulin in addition to K40 are acetylated in various organisms and tissues (Lundby et al., 2012; Weinert et al., 2011; Choudhary et al., 2009; Hansen et al., 2019; Liu et al., 2015a; b). One site is consistently identified in all of these studies: K394. Unlike K40, which is located in the microtubule lumen, K394 sits on the microtubule surface at the αβ-tubulin dimer interface, a region that likely regulates dimer stability and undergoes a conformational change as the dimer is added to the microtubule polymer (Knossow et al., 2020). Based on its position alone, it is possible that K394 could affect dimer stability and microtubule assembly, both of which could in turn affect microtubule stability. Indeed, studies carried out in HeLa cells showed that K394R mutant α-tubulin, which cannot be acetylated, incorporates poorly into microtubules (Liu et al., 2015b). K394 mutations were also uncovered in a screen in CHO cells for α-tubulin mutations that would counteract the stabilising effects of taxol on microtubules (Yin et al., 2013). Thus, K394 acetylation is a compelling candidate to regulate microtubule stability in neurons.

In this study, we asked whether α-tubulin K394 and its acetylation are important to neuronal morphogenesis using the fly neuromuscular junction (NMJ) as a model. We targeted both the K394 residue in endogenous α-tubulin and manipulated a putative K394 deacetylase, HDAC6 (Liu et al., 2015a). These complementary approaches are necessary to (a) determine the importance of K394 to tubulin and microtubule function and (b) interpret the effects of manipulating the modifying enzyme, particularly since the majority of acetyltransferases and deacetylases, including HDAC6, have multiple protein targets. Our studies of the fly NMJ show that K394 is an essential α-tubulin residue that is necessary for axon terminals to develop properly. Mutating K394 results in a dramatic reduction in microtubule stability, and this change in stability underlies the synaptic morphogenesis defects. Combined, our data implicate K394 and its acetylation in regulating microtubule stability during synaptic morphogenesis.

## Results

### Conserved α-tubulin K394 is acetylated in the nervous system and essential for normal synaptic morphogenesis

Acetylation is a post-translational modification that can dynamically modulate protein activity. Several large-scale proteomic studies have revealed that multiple lysines in α-tubulin are acetylated, opening up the possibility that acetylation may regulate microtubule stability and dynamics at sites other than the well-studied K40. One residue is consistently identified in all the protein acetylomes of humans, rats, mice, and flies: K394 (Choudhary et al., 2009; Weinert et al., 2011; Lundby et al., 2012; Liu et al., 2015a; b; Hansen et al., 2019). This conserved residue, K394, is located on the surface of α-tubulin at the αβ-tubulin heterodimer interface (Fig. 1, A and B). We selected this lysine, whose acetylation is conserved across species and which is largely uncharacterised, as a candidate site that may be important for proper microtubule function in neurons.

**Figure 1.**
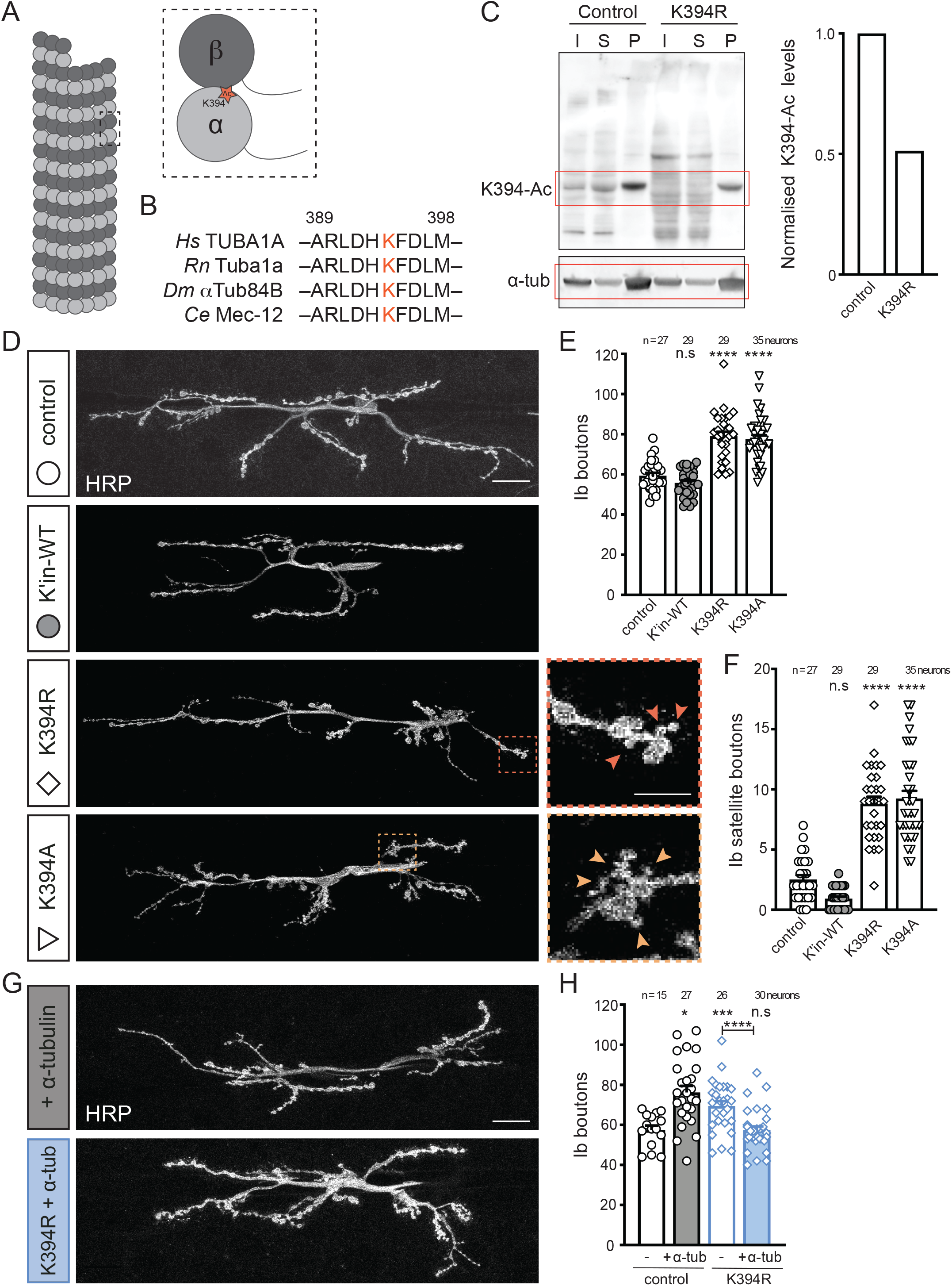
Mutating conserved α-tubulin K394 to block acetylation disrupts synaptic morphogenesis. In this and subsequent figures, all genotypes are homozygous, images are of synaptic terminals at m6/7 in 3^rd^ instar larvae, and quantification is per m6/7, unless otherwise noted. **(A)** Cartoon showing the position of K394 at the αβ-heterodimer interface. **(B)** K394 (red) is conserved between humans, rats, flies and worms. This peptide sequence was used to create the anti-Ac-K394 antibody. ‘ **(C)** Western blot analysis (left) and quantification (right) of K394 acetylation and total α-tubulin levels in control and K394R mutant fly head lysate. The anti-Ac-K394 signal in the tubulin pellet lane is normalised to total α-tubulin and represented as a fraction of the control values. α-tubulin migrates at ∼50 kDa (red box). I=input, S=supernatant, P=pellet. **(D)** Representative images of synaptic terminals in control and mutant animals. K394A mutants die as pupae, and K394R mutants are homozygous viable. Dashed-outline boxes: zoomed-in views to highlight ectopic satellite boutons (arrowheads). Scale bars: 25 µm and 10 µm (dashed-outline boxes). A neuronal membrane marker (HRP) illuminates the synaptic terminals. (E and F) Quantification of total type Ib boutons (E) and satellite boutons (F) in control larvae and larvae in which wild-type (WT) or mutant alleles of tubulin were knocked into endogenous *αTub84B*. **(G and H)** Neuronal over-expression of wild-type α-tubulin (+ α-tubulin) suppresses type Ib bouton overgrowth in K394R mutants. Representative images (G) and quantification (H). *OK6-Gal4* and *UAS-αTub84B*, used to over-express α-tubulin, are hemizygous. Quantification: One-way ANOVA with post-hoc Tukey. All data are mean ± SEM. n.s=non-significant; *p=0.01–0.05; ***p=0.001–0.0001; ****p<0.0001.

To investigate α-tubulin K394, we turned to Drosophila as an in vivo model. As in other organisms, there are multiple α-tubulin isotypes encoded in the fly genome. The four α-tubulin isotypes in Drosophila, which are named based on their cytological positions, are: αTub84B, αTub84D, αTub85E and αTub67C. αTub84B is the essential, predominant, and ubiquitously expressed α-tubulin in flies (Raff, 1984; Yan et al., 2018). αTub84D is almost identical to αTub84B, and it is also ubiquitously expressed; however, its expression is much lower, and αTub84D is not essential for survival (Jenkins et al., 2017). αTub84B is 97% identical to rat α-tubulin, and there are only five non-conservative amino acid differences, the majority of which are in the C-terminal tail. Conservation of the α-tubulin sequence and acetylation, combined with a powerful toolbox for in vivo analysis, makes Drosophila an attractive model system for these studies.

We first set out to determine whether we could detect the acetylation of α-tubulin K394 in the fly nervous system. We were able to successfully raise a peptide antibody against acetylated K394. We pelleted microtubules from fly brain lysate and used the anti-acetylated-K394 antibody to probe a western blot (Fig. 1, B and C). The antibody recognised a prominent band that migrates at the expected size of α-tubulin in the lysate input, supernatant, and microtubule pellet. Next, to determine the specificity of the antibody for the acetylated residue, we probed lysate from the brains of flies expressing α-tubulin with a mutation, K394R, that blocks acetylation (the K394R mutation was knocked into the endogenous αTub84B gene, see below). The antibody signal for acetylated K394 was reduced by at least half, indicating that the antibody recognises the acetylated residue (Fig. 1C). The remaining signal may reflect antibody recognition of other α-tubulin isotypes acetylated at K394, and/or the antibody may retain some affinity for the K394R mutant tubulin. Consistent with published proteomic data showing that K394 is acetylated in the nervous system (Lundby et al., 2012; Liu et al., 2015a), these western blot results indicate that α-tubulin K394 is acetylated in the Drosophila nervous system.

Mutagenesis is a powerful approach to analyse microtubule function in cells. Indeed, the function of α-tubulin K40 acetylation was primarily characterised using K40 mutations before the relevant modifying enzymes were identified decades later (L’Hernault and Rosenbaum, 1985; Hubbert et al., 2002; Akella et al., 2010; Shida et al., 2010). Thus, we leveraged K394 mutations to gain insight into the role that K394 plays in animals and neurons. First, we made a K394A point mutation in the endogenous αTub84B gene. We took advantage of a fly strain that we previously generated that enables us to readily create new alleles of endogenous α-tubulin (Jenkins et al., 2017). The K-to-A mutagenesis revealed that K394 is an essential residue; this is in notable contrast to the non-essential K40, whose acetylation is not required for survival (Jenkins et al., 2017; Kalebic et al., 2013; Kim et al., 2013; Akella et al., 2010). We next made a semi-conservative K394R mutation. Unlike alanine, arginine is similar to lysine in structure and charge, but it cannot be acetylated. Thus, K-to-R mutations are frequently used to mimic the loss of acetylation. While the K394A mutation caused lethality during pupal stages of development, flies homozygous for the K394R mutation were viable. The difference in survival between the K394A and K394R mutations suggests that arginine is likely a sufficient structural substitute for lysine at this position and, thus, K394R is an advantageous mutation for characterising the loss of acetylation at this site.

We then asked whether α-tubulin K394 is important to neuronal morphogenesis. Disrupting microtubule function is frequently associated with defects in neuronal morphology, including the formation of boutons at the fly NMJ (Bodaleo and Gonzalez-Billault, 2016; Penazzi et al., 2016). Boutons are swellings of the axon terminal that house synapses. We used the type Ib (“big”) boutons as a model since they are a well-established system to investigate the role of the cytoskeleton in synaptic growth. We found that both the K394A and K394R mutations resulted in an increase of type Ib boutons and the formation of ectopic small “satellite” boutons (Fig. 1, D-F). This synaptic overgrowth phenotype was suppressed by the over-expression of wild-type α-tubulin specifically in neurons, which suggests that the K394 mutant phenotypes result from a disruption of the presynaptic microtubule cytoskeleton (Fig. 1 G and H). This indicates that α-tubulin K394, which is acetylated in the nervous system, plays a critical role in synaptic morphogenesis.

### Abnormal synaptic morphogenesis in K394R mutant is caused by a decrease in microtubule stability

The morphology phenotype in the K394R mutant animals likely results from a perturbation in tubulin and/or microtubules (Fig. 2A). First, we analysed α-tubulin levels in the K394R mutant. Our western blot analysis of larval brain lysate showed that α-tubulin levels were normal in K394R mutants (Fig. 2B), suggesting that a change in the amount of α-tubulin was unlikely to cause the morphology defect. Next, we tested whether the K394R mutation might affect synaptic morphogenesis by disrupting microtubule stability. We assessed microtubule stability by staining for the microtubule-associated protein Futsch, which is homologous to the mammalian microtubule-associated protein (MAP) MAP1B (Roos et al., 2000; Hummel et al., 2000). Futsch colocalises with stable microtubules, which often form distinctive loops within boutons. The K394R mutation resulted in fewer Futsch-positive loops (Fig. 2, C and D), which indicates a decrease in stable microtubules. We also assessed α-tubulin K40 acetylation, which correlates with stable microtubules in cells (Janke and Magiera, 2020). While anti-acetylated-K40 signal was still present in the K394R mutants, the number of microtubule loops marked by acetylated K40 was reduced (Fig. 2, E and F). This is similar to the Futsch results and consistent with the K394R mutation causing a decrease in microtubule stability. Notably, we found that microtubules were still present even when Futsch levels and Futsch loops were decreased, indicating the K394R mutation did not generally affect microtubule formation (Fig. 2 G). Combined, these results indicate that the K394R mutation does not affect tubulin levels nor drastically decrease microtubule formation; rather, the α-tubulin K394R mutation reduces microtubule stability.

**Figure 2.**
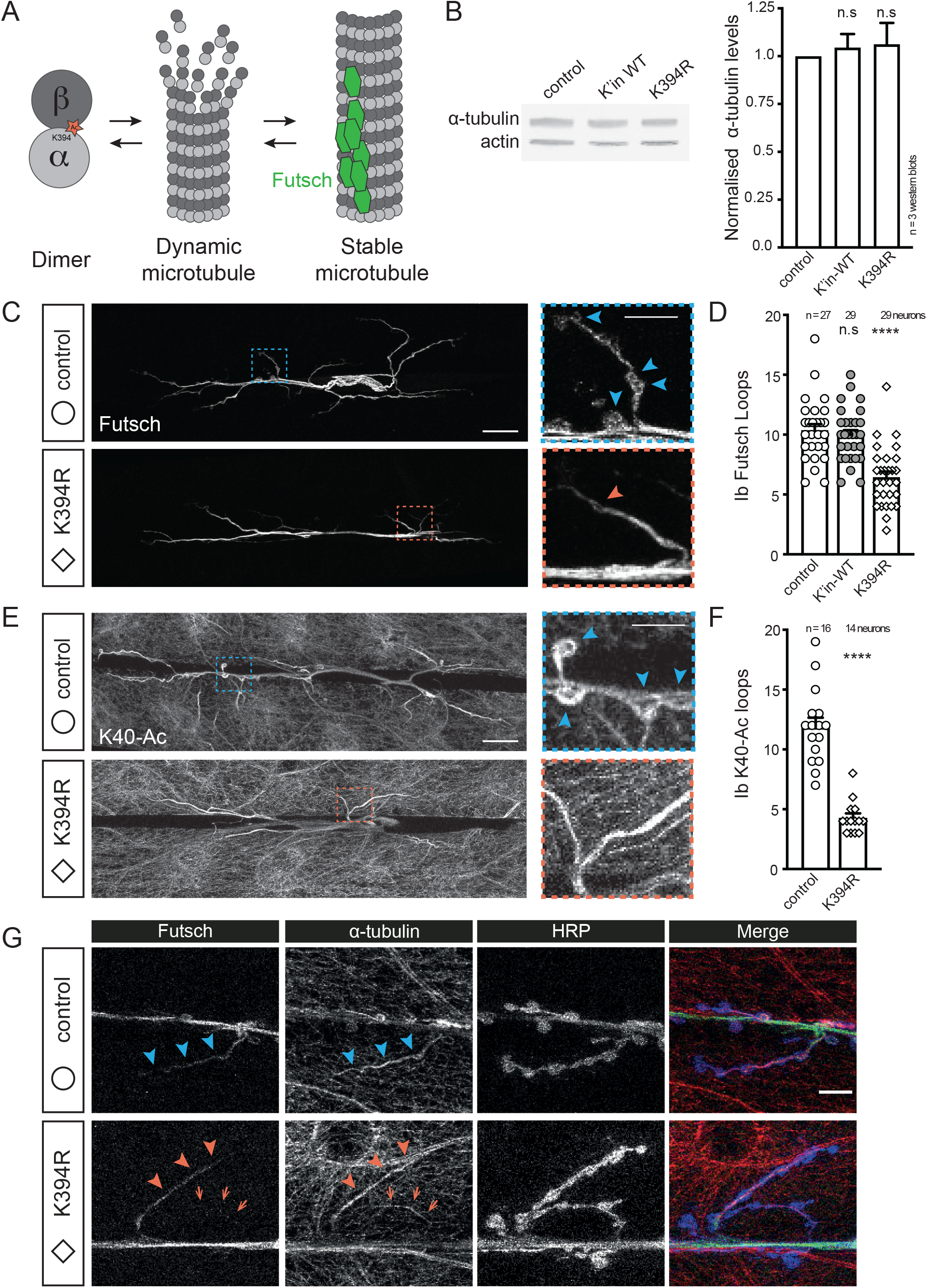
The acetylation-blocking K394R mutation does not affect tubulin levels or microtubule formation but decreases microtubule stability. **(A)** Schematic illustrating free tubulin dimer and two microtubule populations, dynamic and stable. Stable microtubules are bound by Futsch (green hexagons). **(B)** Western blot analysis (top) and quantification (bottom) of α-tubulin levels (DM1α) in control, wild-type αTub84B knock-in, and K394R mutant lysate. DM1α signal is normalised to actin and represented as a fraction of the control values **(C and D)** Representative images (C) of synaptic terminals stained for Futsch, which marks stable microtubules, and the quantification (D) of Futsch-positive microtubule loops (“Futsch loops”) in type Ib boutons in indicated genotypes. Dashed-outline boxes: zoomed-in views to highlight Futsch loops (arrowheads), which are decreased in K394R mutants. Scale bars: 25 µm and 10 µm (dashed-outline boxes). **(E and F)** Representative images (E) of control and K394R synaptic terminals stained for α-tubulin acetylated at K40 (6-11B-1), and the quantification (F) of acetylated-K40-positive microtubule loops (arrowheads) in type Ib boutons. Dashed-outline boxes: zoomed-in views to highlight loops (arrowheads). Scale bars: 25 µm and 10 µm (dashed-outline boxes). **(G)** Decreased Futsch does not reflect a decrease in microtubules in K394R mutants. Larval fillets stained for stable microtubules (Futsch), total α-tubulin (DM1α), and a neuronal membrane marker (HRP). Arrowheads: Futsch loops; arrows: a microtubule branch lacking Futsch. Scale bar: 10 µm. Quantification: One-way ANOVA with post-hoc Tukey (B and D) and Student’s unpaired t-test (F). All data are mean ± SEM. n.s=non-significant; ****p<0.0001.

We next asked whether a decrease in stable microtubules might cause the type Ib synaptic overgrowth phenotype in the α-tubulin K394R mutants. This was a particularly important question to address given that the formation of extra boutons is frequently associated with an increase, not a decrease, in microtubule stability (Bodaleo and Gonzalez-Billault, 2016). Using a combination of pharmacological and genetic approaches, we tested whether enhancing microtubule stability would suppress the formation of additional type Ib boutons in the K394R mutant. First, we treated animals with the microtubule-stabilising drug taxol (also known as paclitaxel). An acute 30-minute treatment of larval fillets with taxol modestly increased the number of Futsch loops in control animals and was sufficient to rescue Futsch loop numbers to normal in the K394R mutant larvae (Fig. 3, A and B). This acute taxol treatment, however, did not suppress the formation of extra boutons in the K394R mutant. Since synaptic boutons develop over several days, it is possible that this 30-minute treatment was too short for taxol to exert an effect on synaptic morphogenesis. We next treated larvae by culturing them for 24 hours on taxol-containing food. The prolonged 24-hour treatment with taxol rescued microtubule stability and also fully suppressed the synaptic overgrowth phenotype in the K394R mutants, reducing the number of boutons in the K394R mutant to normal (Fig. 3, C and D). These taxol experiments indicate that a reduction in microtubule stability likely underlies the overgrowth of type Ib synaptic boutons in the K394R mutant.

**Figure 3.**
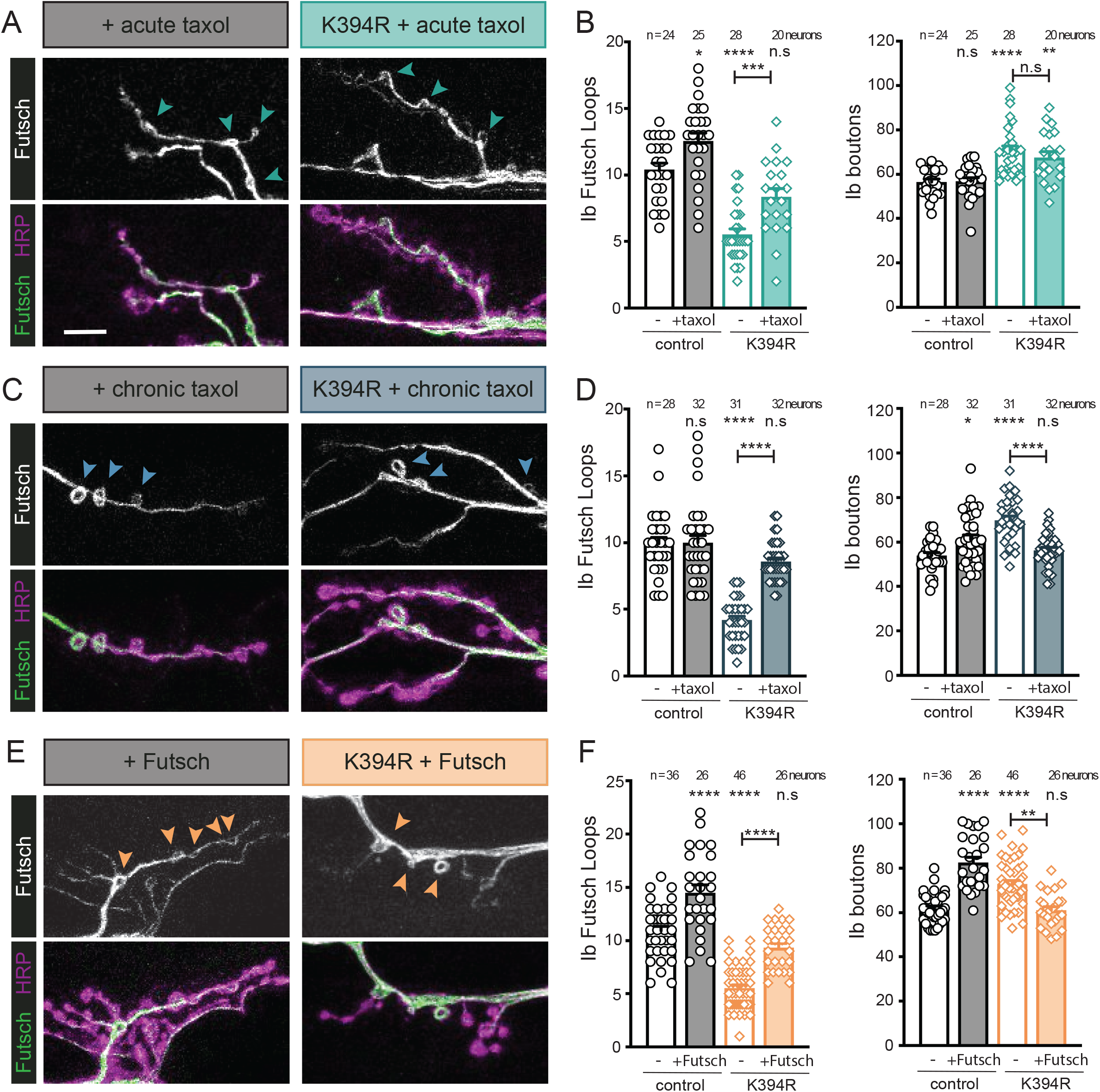
K394R mutant phenotypes are reverted by promoting microtubule stability. Larval fillets stained for stable microtubules (Futsch) and a neuronal membrane marker (HRP). Arrowheads: Futsch loops in type Ib boutons. Futsch loops and type Ib boutons were quantified for each experiment in the indicated genotype and treatment. Scale bar: 10 µm. **(A and B)** Representative images (A) and quantification (B) of dissected control and K394R mutant larvae treated with 50 µM taxol for 30 minutes prior to fixation. **(C and D)** Representative images (C) and quantification (D) of control and K394R mutant larvae raised on food containing 10 µM taxol for 24 hours prior to dissection. **(E and F)** The neuronal over-expression of Futsch affects type Ib bouton growth in control and K394R mutants. *OK6-Gal4* and *UAS-Futsch*, used to over-express Futsch, are hemizygous. Representative images (E) and quantification (F). Quantification: One-way ANOVA with post-hoc Tukey. All data are mean ± SEM. n.s= non-significant; *p=0.01–0.05; **p=0.01–0.001, ***p=0.001–0.0001; ****p<0.0001.

We took an additional genetic approach to test the model that the α-tubulin K394R mutation destabilises microtubules to alter synaptic morphogenesis. We over-expressed the microtubule-stabilising MAP Futsch in control and K394R mutant larvae. In control animals, elevating Futsch levels increased both the number of Futsch loops and type Ib boutons (Fig. 3, E and F), consistent with previous reports that increasing Futsch levels and microtubule stability stimulates type Ib bouton formation (Shi et al., 2019; Roos et al., 2000; Nechipurenko and Broihier, 2012). The over-expression of Futsch in K394R mutant neurons rescued Futsch loop formation, restoring loop numbers to normal, and also suppressed the formation of extra boutons (Fig. 3, E and F). Thus, stabilising microtubules by elevating Futsch levels reverts the K394R mutant phenotypes. Combined with the results of the taxol experiments, these results indicate that the overgrowth of type Ib boutons in the K394R mutant is likely due to a decrease in microtubule stability.

Our results also indicate that the growth of type Ib synaptic boutons depends on achieving a proper level of microtubule stability. We found that the number of type Ib boutons increased when microtubule stability was either reduced by the K394R mutation or enhanced by the over-expression of Futsch (Fig. 3, E and F). Conditions that virtually eliminate stable microtubules, such as in Futsch loss-of-function mutants, reduce type Ib bouton number (Hummel et al., 2000; Roos et al., 2000; Lepicard et al., 2014; Ma et al., 2017). Our data indicate that the K394R mutation significantly diminishes but likely does not eliminate stable microtubules (Fig. 2, C and D). Altogether, our results suggest that type Ib bouton morphogenesis is sensitive to microtubule stability, and the K394R mutation perturbs synaptic growth by decreasing microtubule stability.

### Decreased microtubule stability in K394R mutants is rescued by elevating tubulin cofactor levels

Our data indicate that the K394R mutation decreases microtubule stability. We next sought to determine how the mutation might do so. Previous studies have implicated K394 in microtubule polymerisation and have shown that K394R mutant α-tubulin is not as efficiently incorporated into microtubules as wild-type α-tubulin (Liu et al., 2015b; Szasz et al., 1986, 1993). Based on these findings, we reasoned that the K394R mutation might reduce microtubule stability by disrupting microtubule polymerisation. To test this model, we analysed microtubule dynamics using EB1::GFP, which specifically binds to growing microtubule ends. To eliminate any effects on microtubule growth that might be caused by over-expressing EB1, we tagged endogenous EB1 with the split-GFP peptide sfGFP(11), which can be selectively visualised in neurons that express the complementary peptide sfGFP(1-10) (Fig. 4A) (Kamiyama et al., 2016; Kelliher et al., 2018). Next, since it is difficult to quantitatively analyse microtubule dynamics at the larval NMJ, we turned to sensory neuron dendrites to analyse microtubule polymerisation. If the K394R mutation perturbs microtubule polymerisation, we predicted that EB1::GFP comet number and/or velocity would be reduced. Indeed, we observed significantly fewer EB1::GFP comets in the dendrites of K394R mutant sensory neurons, although there was no significant decrease in comet velocity (Fig. 4, B-E). A reduction in EB1::GFP comets might also reflect an increase in microtubule stability; however, our other data indicate that microtubule stability is decreased, not increased, in K394R mutants. Our data, combined with published reports (Liu et al., 2015b; Szasz et al., 1986, 1993), suggest that the K394R mutation may alter polymerisation such that microtubule stability is reduced.

**Figure 4.**
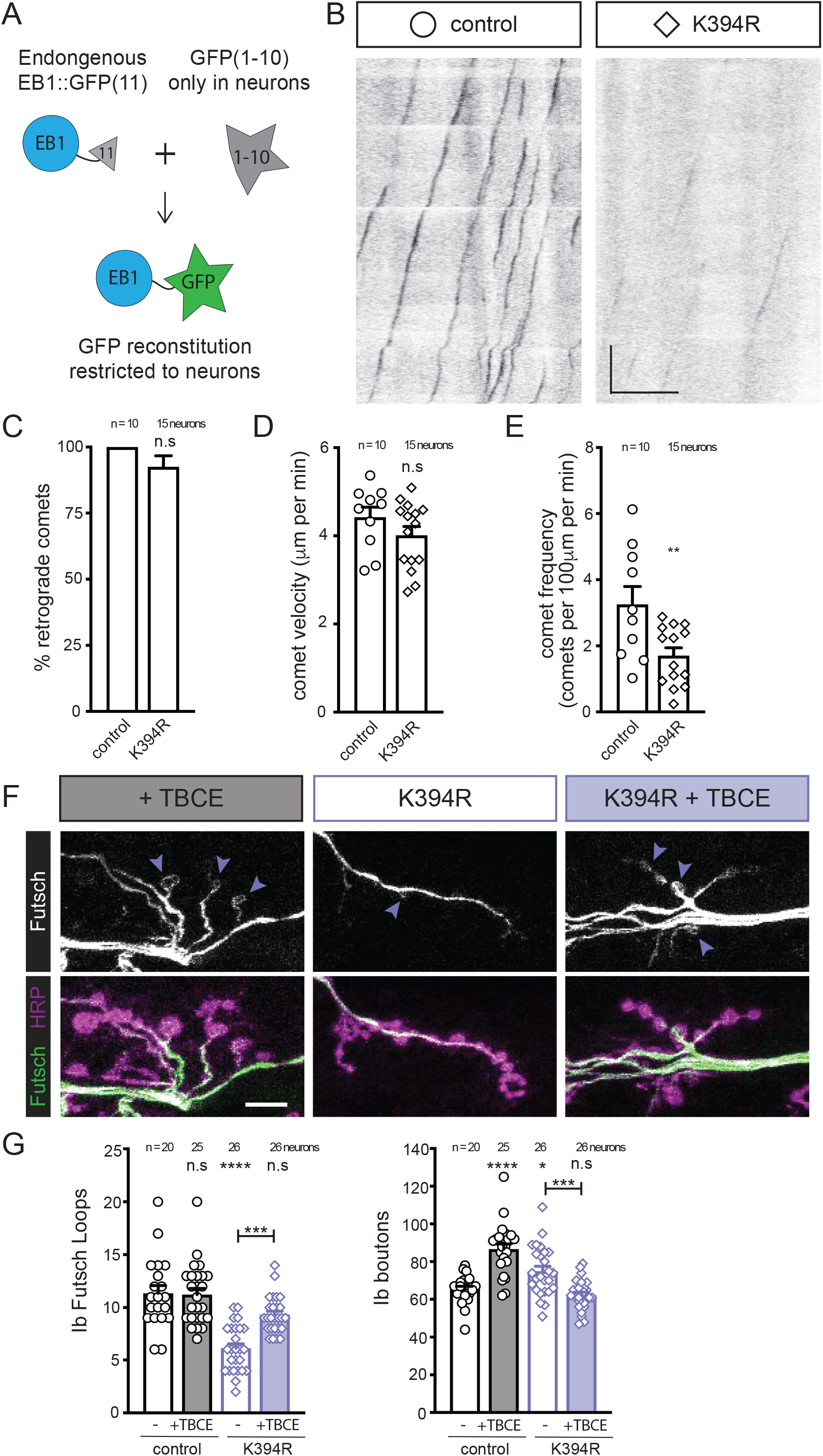
Elevating tubulin chaperone levels reverts K394R mutant phenotypes. **(A)** Cartoon showing the approach to visualise EB1::sfGFP specifically in neurons. Endogenous EB1 was tagged with the split-GFP peptide sfGFP(11), and *pickpocket-Gal4* drove the expression of sfGFP(1-10) in sensory neurons to limit the reconstitution of sfGFP to neurons. **(B)** Representative kymographs of endogenous EB1 dynamics in control and K394R sensory neuron dendrites. Cell body is to the left. *EB1::sfGFP(11), pickpocket-Gal4*, and *UAS-sfGFP(1-10)* are hemizygous. Scale bars: 10 µm (x-axis) and 30 seconds (y-axis). **(C-E)** Quantification of EB1::GFP comet trajectory (C), velocity (D), and frequency (E) in control and K394R mutants. **(F and G)** Representative (F) images and quantification (G) of Futsch loops and type Ib boutons in controls and animals over-expressing TBCE, K394R mutants, and K394R mutants over-expressing TBCE. *OK6-Gal4* and *UAS-TBCE*, used to over-express TBCE, are hemizygous. Larval fillets are stained for stable microtubules (Futsch) and a neuronal membrane marker (HRP). Arrowheads: Futsch loops. Scale bar: 10 µm. Quantification: Student’s unpaired t-test (C-E) and one-way ANOVA with post-hoc Tukey (G). All data are mean ± SEM. n.s=non-significant; *p=0.01–0.05, **p=0.01–0.001, ***p=0.001–0.0001, ****p<0.0001.

Microtubules grow as tubulin dimers are added to microtubule ends. The K394 residue is at the interface of the αβ-tubulin heterodimer. Based on its position, we theorised that K394 may contribute to the stability and/or conformation of the tubulin dimer and, thus, by extension, K394 mutations may affect microtubule stability by perturbing the dimer. Tubulin dimer formation and homeostasis are regulated by a suite of tubulin cofactors, also known as chaperones, and α-tubulin is bound by the tubulin-specific chaperone E (TBCE) (Al-Bassam, 2017). If the K394R mutation perturbs dimer assembly or reduces its stability, this may create a deleterious imbalance in the ratio of dimer to cofactor, e.g., the number of (unstable) dimers may increase relative to TBCE. TBCE and its partner tubulin cofactors likely exist at tightly controlled concentrations at synaptic terminals given that increasing cofactor levels produces synaptic defects (Jin et al., 2009; Okumura et al., 2015). We tested the idea that the K394R mutation perturbs tubulin heterodimer formation or homeostasis by asking whether elevating TBCE expression would reverse the K394R mutant phenotypes. First, as previously reported, we found that higher TBCE levels resulted in significantly more boutons than controls (Jin et al., 2009), albeit the number of Futsch loops was unaffected (Fig. 4, F and G). Increasing TBCE levels in the K394R mutant suppressed the formation of extra boutons and significantly, also restored the formation of Futsch loops to wild-type levels (Fig. 4, F and G). Thus, the effects of the K394R mutation on microtubule stability can be reversed by increasing the amount of TBCE. While this experiment does not directly test the effects of the K394R mutation on tubulin dimer stability, these results are consistent with the notion that the K394R mutation may destabilise or otherwise alter the tubulin dimer such that microtubule stability is reduced.

### α-tubulin K394 acetylation is regulated by HDAC6

We next wanted to identify the enzyme(s) that regulate the acetylation of K394. A previous study using mass spectrometry revealed that the deacetylase HDAC6 targets several tubulin residues, including both α-tubulin K40 and K394 (Liu et al., 2015a). To determine whether HDAC6 indeed regulates the acetylation of α-tubulin at K394, we used our anti-Ac-K394 antibody to probe microtubules pelleted from the lysate of control and HDAC6-knock-out fly brains. If HDAC6 is responsible for deacetylating K394, K394 acetylation should increase when HDAC6 is knocked-out. When we pelleted microtubules from HDAC6 knock-out brain lysate, we indeed saw a dramatic increase in K394 acetylation (Fig. 5A). This indicates that HDAC6 promotes the deacetylation of K394, either directly or indirectly.

**Figure 5.**
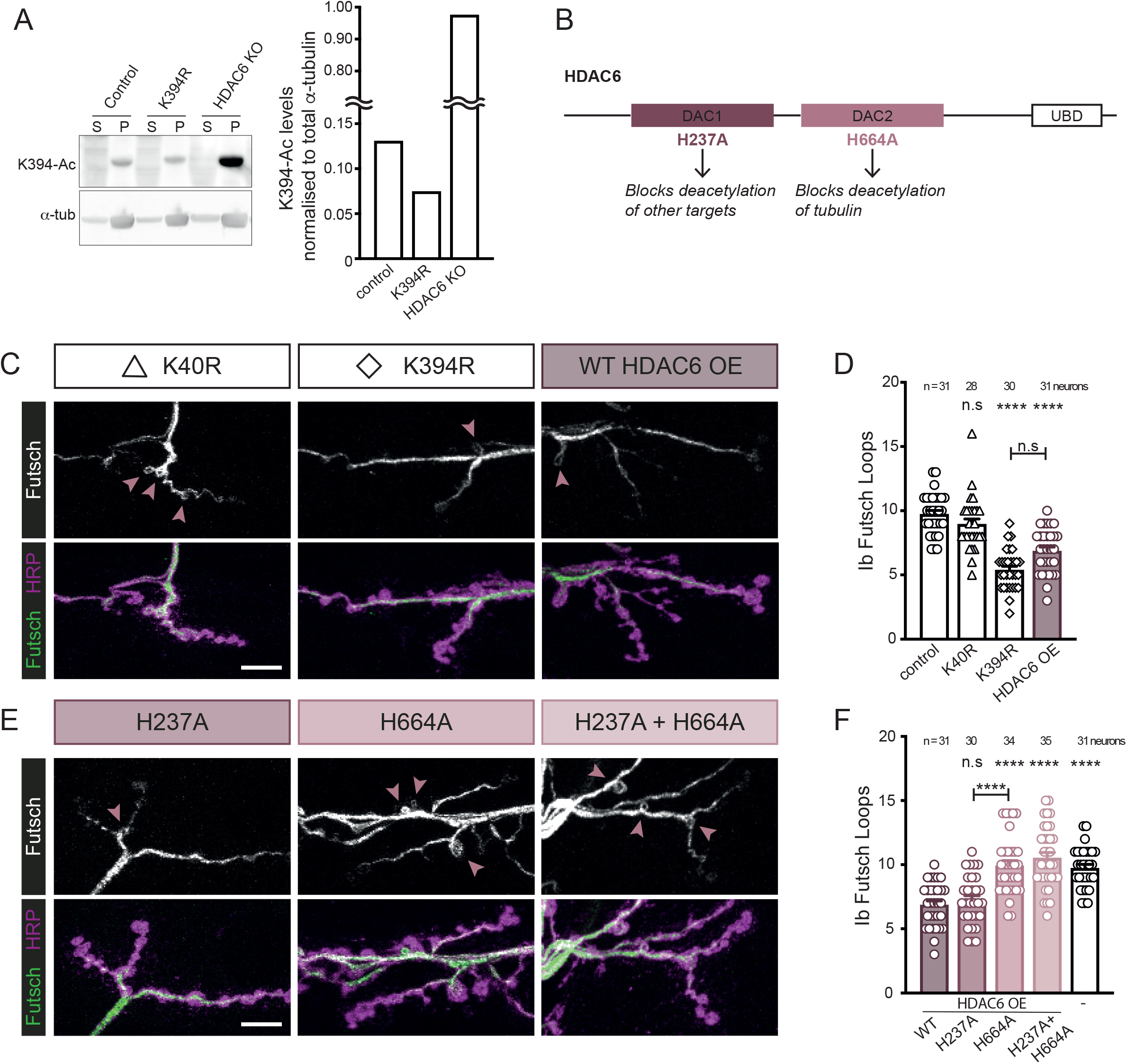
α-tubulin K394 acetylation is regulated by HDAC6. **(A)** Western blot analysis (left) and quantification (right) of K394 acetylation of microtubules pelleted from control, K394R, and HDAC6-knockout (KO) fly heads. The anti-Ac-K394 signal in the tubulin pellet lane is normalised to total α-tubulin. S=supernatant, P=pellet. **(B)** Cartoon illustrating HDAC6 and its key domains, including two deacetylase domains (DAC1 and DAC2) and a ubiquitin-binding domain (UBD). The functional outcomes of point mutations in DAC1 and DAC2 are indicated. **(C-F)** The activity of DAC2 is necessary to reduce microtubule stability similar to the K394R mutants. Representative images (C and E) and quantification (D and F) of Futsch loops and type Ib boutons in control, K40R and K394R mutants, and animals over-expressing wild-type HDAC6 (HDAC6 OE) or mutant HDAC6 (H237A and/or H664A). All genotypes were included in the experiment, whose quantification is split between panels D and F. The quantification of the control and animals over-expressing wild-type HDAC6 are included in both D and F. *OK6-Gal4* and *UAS-HDAC6* (wild-type and mutant constructs), used to over-express HDAC6, are hemizygous. Larval fillets are stained for stable microtubules (Futsch) and a neuronal membrane marker (HRP). Arrowheads: Futsch loops. Scale bar: 10 µm. Quantification: One-way ANOVA with post-hoc Tukey. All data are mean ± SEM. n.s=non-significant; *p=0.01–0.05; ***p<0.0001.

Next, we leveraged HDAC6 to ask whether K394 acetylation is important to synaptic microtubule stability. Since K394R is an acetylation-blocking mutation, we decided to over-express HDAC6, which should reduce acetylation, and compare the two. When we over-expressed wild-type HDAC6, we found that it reduced microtubule stability similar to the acetylation-deficient K394R mutant (Fig. 5, B-D). HDAC6 is a multi-functional enzyme that has a role in diverse cellular activities: it targets multiple sites in tubulin and other proteins for deacetylation, and HDAC6 also functions via its ubiquitin-binding domain (Valenzuela-Fernández et al., 2008). Therefore, we took advantage of known mutations in HDAC6 that selectively abrogate its deacetylation activity. HDAC6 has two deacetylase domains (DAC1 and DAC2), only one of which, the second domain (DAC2), acts on tubulin (Fig. 5B) (Xiong et al., 2013; Haggarty et al., 2003; Kaluza et al., 2011). We focused on the effects of HDAC6 on microtubule stability as DAC1 has been implicated in disrupting synaptic morphogenesis by acetylating Bruchpilot, a scaffolding component of the active zone (Miskiewicz et al., 2014). We over-expressed mutant HDAC6 enzymes that were defective in deacetylating non-tubulin targets (H237A in DAC1), tubulin (H664A in DAC2), or both (H237A, H664A double mutant). Over-expression of the mutant HDAC6 defective in deacetylating non-tubulin targets (H237A) reduced stable microtubules similar to the over-expression of wild-type HDAC6 (Fig. 5, E and F). In contrast, the over-expression of HDAC6 mutants that could not deacetylate tubulin (H664A and the H237A, H664 double mutant) had no effect on stable microtubules (Fig. 5, E and F). Thus, HDAC6 affects microtubule stability in type Ib boutons via the domain that deacetylates tubulin (DAC2). Notably, mutating α-tubulin K40, another HDAC6 target, did not affect the number of Futsch loops, indicating that HDAC6 was unlikely to exert its effects on microtubule stability in type Ib boutons by modulating K40 acetylation (Fig. 5 C). Combined, our results suggest that acetylation of α-tubulin K394 is likely regulated by HDAC6, and this is important to microtubule stability during the morphogenesis of type Ib synaptic terminals.

### The K394R mutation and alterations in microtubule stability have neuron-type-specific effects on synaptic morphogenesis

We were struck by the observation that type Ib synaptic growth increased when microtubule stability was either reduced, as in the K394R mutation, or enhanced, as by the over-expression of Futsch in control animals. Type Ib synaptic terminals are known to be both physiologically and morphologically plastic (Aponte-Santiago and Littleton, 2020). For example, in response to enhanced activity, type Ib neurons generate additional boutons, the growth of which depends on the microtubule cytoskeleton (McLaughlin et al., 2016; Coombes et al., 2020). We considered the possibility that the growth of the type Ib synaptic terminals in response to any alteration in microtubule stability may be characteristic of this particular neuron type. Another motoneuron type that innervates the same m6/7 muscle pair, type Is (“small”), is functionally and morphologically distinct from type Ib, and, unlike the type Ib neurons, type Is neurons do not form ectopic boutons in response to changes in activity (Fig. 6A) (Aponte-Santiago and Littleton, 2020). Given these differences, we asked whether the type Is neurons would respond differently than type Ib to changes in microtubule stability.

**Figure 6.**
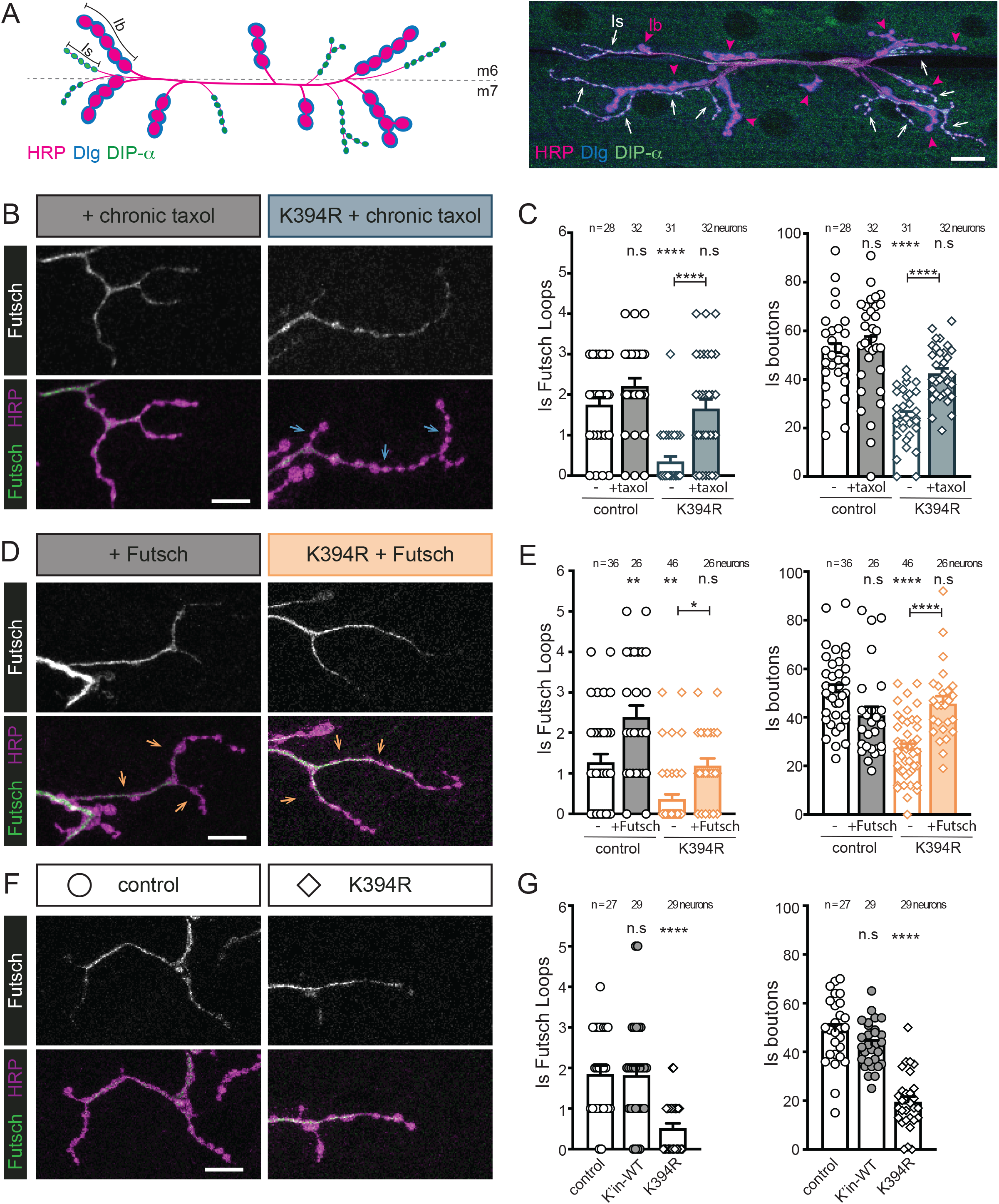
Manipulation of microtubule stability has neuron-type-specific effects. **(A)** Cartoon (left) and representative image (right) of type Ib and Is boutons that synapse on muscles 6 and 7 (m6 and m7) visualised with a neuronal membrane marker (HRP), Discs large 1 (Dlg), and *DIP-α-GFP*. The type Ib and Is boutons can be distinguished by differences in size (Ib are “big,” and Is are “small”) and molecular make-up: the type Ib boutons have high levels of Dlg and type Is neurons express *DIP-α-GFP*. Scale bar: 20 µm. **(B-G)** Larval fillets stained for stable microtubules (Futsch) and a neuronal membrane marker (HRP). Arrows indicate branches of type Is boutons. Futsch loops and type Is boutons were quantified. Scale bar: 10 µm. **(B and C)** Representative images (B) and quantification (C) of control and K394R mutant larvae raised on food containing 10 µM taxol for 24 hours prior to dissection. **(D and E)** Effects of Futsch over-expression on microtubule stability and the growth of type Is boutons. Representative images (D) and quantification (E) of control and neurons over-expressing Futsch (*OK6-Gal4* and *UAS-Futsch*, hemizygous). **(F and G)** Representative images (F) and quantification (G) of control and K394R mutant. Quantification: One-way ANOVA with post-hoc Tukey. All data are mean ± SEM. n.s=non-significant; ***p=0.01–0.001; ****p<0.0001.

We manipulated microtubule stability in several ways to test the response of type Is neurons. First, we asked whether enhancing microtubule stability in control animals would increase the number of type Is boutons. As we had done previously, we enhanced microtubule stability by raising control larvae on taxol-containing food or by over-expressing Futsch. Whereas both these manipulations increased type Ib bouton number (Fig. 3, C, D, E, and F), type Is bouton number was unchanged by either of these microtubule-stabilising treatments (Fig. 6, B-E). Thus, unlike type Ib neurons, type Is neurons do not generate additional boutons when microtubule stability is enhanced. Next, we investigated the effects of the microtubule destabilising mutation K394R on type Is bouton growth. Consistent with its effects on the type Ib cytoskeleton, the K394R mutation decreased the number of Futsch loops in type Is synaptic terminals (Fig. 6, F and G). However, the K394R mutation resulted in significantly fewer type Is boutons than controls (Fig. Fig. 6, F and G). This decrease in type Is bouton number was rescued by treatment with taxol or by elevating Futsch levels (Fig. 6, B-E), which indicated that type Is bouton loss was due to decreased microtubule stability. These results surprisingly reveal that the K394R mutation generates opposite synaptic growth phenotypes in two different neuron types even though the mutation has the same effect on microtubule stability. Combined with our other results, this indicates that the synaptic growth effects induced by altering microtubule stability depend on the cellular context that varies between different neuron types.

## Discussion

Microtubules are integral to neuronal structure and activity, and their function can be regulated by post-translational modifications such as acetylation. While the catalogue of tubulin acetylation sites has grown since tubulin was first shown to be acetylated over thirty-five years ago (L’Hernault and Rosenbaum, 1985), a central challenge remains: to understand how acetylation regulates microtubule function in neurons and other cell types. Here, we investigated a conserved acetylation site in α-tubulin, K394, that has been consistently identified in acetylome studies, yet remained uncharacterised. Leveraging an in vivo fly model system and mutagenesis of endogenous α-tubulin, we discovered that K394 is an essential residue that is acetylated in the nervous system and is required for microtubule stability and proper synaptic terminal development. Acetylation of K394 increases when the deacetylase HDAC6 is reduced, indicating that HDAC6 regulates K394 acetylation. Consistent with HDAC6 functioning to deacetylate K394 at developing synaptic terminals, we found that the over-expression of HDAC6 destabilises microtubules similar to the acetylation-blocking K394R mutation, and this activity depends on its tubulin deacetylase domain. Together, our data suggest a model in which the α-tubulin K394 and its acetylation are important for the stability of microtubules during synaptic terminal development.

Our mutagenesis revealed that reduced microtubule stability in K394R mutants leads to striking neuron-type-specific morphogenesis defects. The type Ib and Is neurons have distinct roles in larval locomotion and respond differently to changes in neurotransmission (Aponte-Santiago and Littleton, 2020). For example, type Ib synaptic terminals can increase their number of boutons in response to changes in activity whereas type Is do not. Our work indicates that these two neuron types also respond differently to cytoskeletal changes: type Ib neurons overgrow when microtubule stability is reduced or increased. In contrast, the type Is neurons are sensitive to reduced microtubule stability, but enhanced microtubule stability does not induce ectopic synaptic growth. Recent studies have uncovered some crosstalk between type Ib and Is when the activity (or presence) of one is disrupted, and it is possible that this may contribute to some of the morphological differences we observe (Wang et al., 2021; Aponte-Santiago et al., 2020) Our findings raise the possibility that the microtubule cytoskeleton may be regulated in a neuron-type-specific manner in order to create the discrete morphologies that underlie the distinct functions of type Ib and Is neurons. It is likely that differences in the expression or activity of cytoskeletal regulators between these neuron types result in divergent phenotypes when microtubule stability is altered. Neuron-type-specific suites of cytoskeletal regulators could contribute to the intrinsic differences between these two neuron types.

Strikingly, the K394R mutant phenotype can be reversed by increasing the levels of the α-tubulin cofactor TBCE, which has been directly implicated in both the assembly and homeostasis of αβ-tubulin dimers (Al-Bassam, 2017). Reducing TBCE levels reduces stable microtubules, which suggests that defects in either αβ-tubulin dimer formation or homeostasis ultimately affect microtubule stability (Jin et al., 2009). TBCE functions as part of a cofactor complex that is postulated to regulate the interaction between α- and β-tubulin at the heterodimer interface, which is where K394 is located (Nithianantham et al., 2015; Serna et al., 2015). K394 belongs to a patch of three basic residues at the heterodimer interface that have been postulated to regulate polymerisation (Szasz et al., 1986). Although the over-expression of TBCE results in microtubule loss in some cell types, such as HeLa cells (Bhamidipati et al., 2000), elevating TBCE levels has also been reported to enhance microtubule stability at the fly neuromuscular junction (Jin et al., 2009). In our experiments, the over-expression of TBCE does not have a significant effect on stable microtubules, although it induces the formation of extra boutons, consistent with a previous report (Jin et al., 2009). Both the position of K394 at the αβ-tubulin heterodimer interface and the ability of TBCE over-expression to reverse K394R mutant phenotypes suggests that the mutation disrupts αβ-tubulin dimer stability and/or homeostasis. Thus, one likely model is that the K394R mutation reduces microtubule stability through an effect on the dimer. This in turn suggests that K394 acetylation might similarly regulate microtubule stability by affecting αβ-tubulin heterodimer formation and/or homeostasis. Interestingly, the acetylation of a β-tubulin residue (K252) that also resides at the αβ-tubulin heterodimer interface is proposed to regulate microtubule polymerisation by affecting dimer conformation (Chu et al., 2011). Like β-tubulin K252 acetylation, it is possible that acetylation of α-tubulin K394 may impinge on the αβ-tubulin heterodimer to regulate microtubule stability.

Elucidating the cellular effects of a microtubule modification includes identifying and manipulating the relevant modifying enzymes. Our finding that K394 acetylation dramatically increases in the HDAC6 knock-out indicates that the deacetylase negatively regulates K394 acetylation, consistent with a previous report (Liu et al., 2015a). While additional experiments are needed to determine whether HDAC6 directly deacetylates K394, our findings indicate that the enzyme regulates K394 acetylation in neurons in vivo. HDAC6 targets other sites in α-tubulin, including the well-studied K40 (Liu et al., 2015a). α-tubulin K40 acetylation has long been correlated with stable microtubules, and knocking-out HDAC6 increases K40 acetylation, which, in turn, is frequently associated with an increase in microtubule stability (Janke and Magiera, 2020). For example, the loss of HDAC6 in fly and mouse muscle cells makes microtubules more stable and resistant to either cold or nocodazole treatment (Mao et al., 2017; Osseni et al., 2020). While such examples of HDAC6’s effects on microtubule stability have been linked to K40 acetylation, our results suggest that some of these effects might also be due to HDAC6 regulating the acetylation of K394. We found that HDAC6 over-expression reduced microtubule stability in synaptic terminals similar to blocking acetylation at K394 but not K40 (K394R and K40R mutants, respectively). In addition, we observed that K40 is still acetylated in K394R mutants, which indicates that the effects of K394R are not likely due to a reduction in K40 acetylation. Thus, our K394 mutagenesis and HDAC6 experiments, combined with published quantitative mass spectrometry results (Liu et al., 2015a), suggest that HDAC6 regulates α-tubulin K394 acetylation to modulate microtubule stability in developing synaptic boutons.

Ultimately it will be important to identify the enzyme that acetylates α-tubulin K394. The well-studied α-tubulin K40 is acetylated by α-tubulin acetyltransferase (αTAT), but some evidence suggests that αTAT does not target K394. The mammalian enzyme αTAT1 is found predominantly in the microtubule lumen where K40 is positioned but has minimal interaction with the microtubule surface where K394 is positioned, which is inconsistent with K394 being an αTAT substrate (Coombes et al., 2016; Howes et al., 2014; Szyk et al., 2014). Moreover, we and others have previously shown that the loss of αTAT in Drosophila does not phenocopy the K394R mutant: in αTAT knock-out animals, microtubule dynamics are increased, and microtubule stability is unchanged at synaptic terminals (Coombes et al., 2020; Yan et al., 2018; Topalidou et al., 2012; Neumann and Hilliard, 2014; Wei et al., 2017). Nonetheless, we are currently pursuing αTAT and additional candidates to identify the acetyltransferase that targets α-tubulin K394. Although we do not yet know the relevant acetyltransferase, our data suggest a model in which acetylation at K394 has a role in regulating microtubule stability during neuronal development.

## Acknowledgements

We thank the Bloomington Drosophila Stock Center for fly strains and Dr. Richard Ordway (Penn State University) and the Developmental Studies Hybridoma Bank for antibodies. We thank members of the O’Connor-Giles laboratory (Brown University) and Gardner laboratory (University of Minnesota) for helpful advice and input, and members of the Wildonger laboratory for thoughtful discussions and comments on the manuscript. This work was generously supported by the National Institutes of Health (NIH) grant R01NS116373 to J.W.

## Author contributions

H.A.J.S., B.V.J, and J.W. conceived of the study. H.A.J.S., D.J.S., and P.J.V. conducted experiments and analysed data. B.V.J., H.A.J.S., P.J.V., and D.J.S. designed and generated the α-tubulin alleles; S.Z.Y. and D.J.S. designed and generated the EB1::GFP fly strain. H.A.J.S. and J.W. wrote the manuscript with input from all the authors.

## Materials and Methods

### Fly Husbandry and Strains

All stocks and crosses were maintained at 25°C under 12-hour light-dark cycles on standard cornmeal-molasses food unless otherwise stated. The following fly strains are from the Bloomington Drosophila Stock Center (BDSC): *w*^*1118*^ (stock # 6326), *Futsch*^*EP1419*^ (stock # 10751), *OK6-Gal4* (stock # 64199), *pickpocket-Gal4* (stock # 320790), *UAS-TBCE* (stock # 34536), *UAS-HDAC6*^*WT*^ (stock # 51181), *UAS-HDAC6*^*H664A*^ (stock # 51184), *UAS-HDAC6*^*H237A*^(stock # 51183) and *UAS-HDAC6*^*H237A+H664A*^ (stock # 51185). *UAS-sfGFP(1-10)* was created as previously described (Kamiyama et al., 2016; Kelliher et al., 2018).

All *αTub84B* knock-in strains were made using *αTub84B*^*attP-KO*^, in which the αTub84B gene has been replaced with an *attP* site for easy knock in of new tubulin alleles (Jenkins et al., 2017). To create new *αTub84B* alleles, the *pGE-attB-GMR* integration plasmid containing the new allele was injected into *αTub84B*^*attP-KO*^ embryos expressing the integrase PhiC31 (BestGene Inc.). The following mutations were introduced into the integration plasmid by Phusion high fidelity polymerase: *K394R* and *K394A. αTub84B*^*K’in-WT*^ and *αTub84B*^*K40R*^ were also generated using this method (Jenkins et al., 2017). *UAS-αTub84B* was made by cloning *αTub84B* into the *pIHEU-MCS* plasmid (# 58375; Addgene) and was injected into *P{CaryP}attP2* (BestGene Inc.).

CRISPR-Cas9-mediated genome editing was used to create *EB1::sfGFP(11)*. A plasmid to express a guide RNA targeting the 3’-end of the *EB1* coding sequence (*pBSK-U63-EB1-gRNA*) was injected with a repair template into *nos-Cas9* embryos (BDSC stock # 78782) (BestGene Inc.). The repair template encoded a linker (GGSGG) plus seven tandem copies of sfGFP(11) separated by GGSGG linkers. To identify integration events, the repair template also included a *3xP3-dsRed* cassette, which was flanked by the *piggyBac* inverted terminal repeat sequences. Candidate progeny expressing *3xP3-DsRed* were crossed to a transposase-expressing strain to scarlessly excise *3xP3-dsRed* cassette. The EB1::sfGFP(11) gene was then fully sequenced. EB1 guide RNA target sequence: 5’-TAATACTCCTCGTCCTCTGG-3’.

Linker sequence: 5’-GGCGGATCCGGCGGA-3’.

sfGFP(11)x7: 5’-

CGTGACCACATGGTCCTTCATGAGTATGTAAATGCTGCTGGGATTACAGGTGGCTCTGGAGGTAGAGATCATATGGTTCTCCACGAATACGTTAACGCCGCAGGCATCACTGGCGGTAGTGGAGGACGCGACCATATGGTACTACATGAATATGTCAATGCAGCCGGAATAACCGGAGGGT CCGGAGGCCGGGATCACATGGTGCTGCATGAGTATGTGAACGCGGCGGGTATAACTGGTGGGTCGGGCGGACGAGACCATATGGTGCTTCACGAATACGTAAACGCAGCTGGCATTACTGGCGGATCAGGTGGCAGGGATCACATGGTACTCCATGAGTACGTGAACGCTGCTGGAATCACAGGCGGTAGCGGCGGTCGGGACCATATGGTCCTGCACGAATATGTCAATGCTGCCGGTATCACC-3’

### Imaging and Immunohistochemistry

All imaging was done on an SP5 confocal microscope (Leica Microsystems) using a 40 × 1.3 NA oil-immersion objective. Muscle pair 6/7 in abdominal segment A2 were imaged and analysed unless otherwise noted.

Wandering third instar larvae were dissected in PHEM buffer (80 mM PIPES pH 6.9, 25 mM HEPES pH 7.0, 7 mM MgCl_2_, 1 mM EGTA) and fixed in 4% PFA in 1X phosphate-buffered saline (PBS) with 3.2% sucrose for 30 minutes (mins), permeabilised in 1XPBS with 0.3% Triton-X100 for 20 mins, quenched in 50 mM NH_4_Cl for 10 mins, blocked in buffer containing 2.5% BSA (catalogue number A9647, Sigma), 0.25% FSG (catalogue number G7765, Sigma), 10 mM glycine, 50 mM NH_4_Cl, 0.05% Triton-X100 for at least 1 hour at room temperature. Fillets were then incubated in primary antibody in blocking buffer overnight at 4°C, washed in 1XPBS with 0.1% Triton-X100 at room temperature and incubated with secondary antibody in blocking buffer overnight at 4°C in the dark. After washing in 1XPBS with 0.1% Triton-X100, fillets were mounted in elvanol containing antifade (polyvinyl alcohol, Tris 8.5, glycerol and DABCO; DABCO, catalogue number 11247100, Fisher Scientific, Hampton, NH). The following antibodies were used on dissected larval fillets: mouse anti-Futsch 22C10 (1:50, Developmental Studies Hybridoma Bank, Iowa City, IA), rabbit anti-Futsch-LC (1:5,000, gift of R. Ordway, Penn State University)(Zou et al., 2008), mouse-anti-Dlg (1:100, Developmental Studies Hybridoma Bank, Iowa City, IA), mouse anti-α-tubulin DM1α (1:500, Sigma-Aldrich), mouse anti-acetylated-K40 6-11B-1 (1:500, Sigma-Aldrich), goat anti-HRP conjugated Alexa Fluor 647 (1:3000, or 0.5 µg mL^-1^, Jackson ImmunoResearch, West Grove, PA), Dylight 550 anti-mouse (1:1000, or 0.5 µg mL^-1^, ThermoFisher, Waltham, MA), Dylight 488 anti-mouse (1:1000, or 0.5 µg mL^-1^, ThermoFisher, Waltham, MA).

### Taxol Treatment

For acute taxol treatment, wandering third instar larvae were dissected in PHEM buffer containing 50 μM taxol and incubated for 30 mins. Subsequently, larvae were fixed and stained following the methods above. For chronic taxol treatment, 24 hours before dissections, larvae were transferred onto cornmeal molasses food containing 10 μM taxol. Larvae were then dissected following the methods above.

### Quantification of Bouton Number and Futsch and acetylated-K40 Loops

Type Ib and Is boutons and Futsch and acetylated-K40 loops in these boutons were quantified at m6/7 of segment A2 in blinded images. Satellite boutons were defined as five or fewer smaller sized boutons coming from the main terminal branch of the nerve. Futsch and acetylated-K40 loops were defined as complete, unbroken loops of signal within a bouton.

### EB1::GFP Imaging and Analysis

Wandering third instar larvae were immobilised in 50% glycerol in 1XPBS on a slide between two strips of vacuum grease. Dorsal class IV ddaC sensory neurons in abdominal segments 2-4 were imaged using a 40 × 1.3 NA oil-immersion objective. EB1 comet trajectories were visualised by reconstituting GFP fluorescence specifically in class IV neurons: *pickpocket-Gal4* was used to drive the expression of *UAS-sfGFP(1-10)* in *EB1::sfGFP(11)* larvae. Videos were captured at a rate of 1.358 frames/second and a resolution of 1,024 x 512 pixels. Videos were stabilised using the FIJI Image Stabilizer plugin and kymographs generated in FIJI. Comet direction, velocity and frequency were then analysed in MetaMorph and data exported to Excel for analysis.

### Anti-Acetylated-K394 Custom Antibody

To detect acetylated α-tubulin K394, the following peptide was used to generate polyclonal antibodies in rabbits: ARLDH(AcK)FDLMYAK (ThermoFisher Scientific). The antibodies were purified via negative selection against non-acetylated peptide ARLDHKFDLMYAK (ThermoFisher Scientific).

### Analysis of α-tubulin Levels and Microtubule Pelleting

To analyse α-tubulin levels, 10 larval fly brains were dissected in 1XPBS, then 30 µL of 1XSDS loading buffer was added, and the sample was boiled for 10 mins. Lysate from the equivalent of 2 brains (6 µL) was loaded per lane.

Microtubules were pelleted from the lysate of fly heads from controls (*w*^*1118*^) and K394R mutants. Flies were flash frozen then vortexed for 1 min, and a sieve was used to separate heads from bodies. Heads were homogenised in 1 mL BRB-80 lysis buffer (80 mM PIPES, 1 mM MgCl_2_, 1 mM EGTA, 1 mM DTT, cOmplete EDTA-free protease inhibitor cocktail [Roche], 100 nM TSA, 1 mM sodium butyrate, 10 mM nicotinamide, 2 µM SAHA) per 1 gram of heads. To remove cell debris, samples were centrifuged at 20,000g at 4°C for 30-45 mins. Lysate was then further clarified by centrifuging at 209,000g for 14 mins at 4°C. 3 mM GTP and 20 µM taxol was then added to the lysate and microtubules allowed to polymerise for 30 mins at room temperature followed by incubation for 30 mins on ice. Samples were then centrifuged through a 15% sucrose cushion at 45,000g for 4 mins at 4°C to pellet microtubules. Pellets were resuspended in BRB-80 lysis buffer with 20 µM taxol.

### Immunoblotting

Samples were run on an SDS-PAGE gel, and then proteins were transferred to low-fluorescence PVDF membrane (catalogue number GE10600022, GE Health and Life Sciences). Membranes were blocked in 5% milk in 1X Tris-buffered saline (TBS) with 0.1% Tween-20 (0.1% TBST) for 1 hour at room temperature and incubated with primary antibody in block overnight at 4°C. After washing in 0.1% TBST membranes were incubated with secondary antibodies for 2-4 hours at room temperature. Membranes were then imaged using either chemiluminescence (Super Signal West Femto ECL, ThermoFisher Scientific) or fluorescence. The following antibodies were used for western blotting: mouse anti-α-tubulin DM1α (1:1000, Sigma-Aldrich), mouse anti-actin C4 (1:1000, Sigma-Aldrich), rabbit anti-Ac-K394 (1:50, this study), Cy5 anti-mouse (1:10000, Jackson ImmunoResearch, West Grove, PA), and HRP-conjugated anti-mouse (1:10000, Bio-Rad Laboratories).

### Statistical Analysis

Data were blinded prior to analysis. Statistical analysis was performed in Excel and GraphPad Prism using a significance level of p < 0.05. Data were first analysed for normality using the Shapiro-Wilk test. Normally distributed data was then analysed for equal variance and significance using either an F-test and Student’s unpaired t-test (two samples) or one-way ANOVA with post-hoc Tukey (multiple samples). Data sets that were not normally distributed were analysed using Mann-Whitney U test (two samples) or Kruskal-Wallis test with post-hoc Dunn test for significance (multiple samples).

## Notes

### Competing Interest Statement

The authors have declared no competing interest.

## References

Akella, J.S., D. Wloga, J. Kim, N.G. Starostina, S. Lyons-Abbott, N.S. Morrissette, S.T. Dougan, E.T. Kipreos, and J. Gaertig. 2010. MEC-17 is an α-tubulin acetyltransferase. Nature. 467:218–222. doi:10.1038/nature09324.

Al-Bassam, J. 2017. Revisiting the tubulin cofactors and Arl2 in the regulation of soluble αβ-tubulin pools and their effect on microtubule dynamics. Mol Biol Cell. 28:359–363. doi:10.1091/mbc.e15-10-0694.

Aponte-Santiago, N.A., and J.T. Littleton. 2020. Synaptic Properties and Plasticity Mechanisms of Invertebrate Tonic and Phasic Neurons. Front Physiol. 11:611982. doi:10.3389/fphys.2020.611982.

Aponte-Santiago, N.A., K.G. Ormerod, Y. Akbergenova, and J.T. Littleton. 2020. Synaptic plasticity induced by differential manipulation of tonic and phasic motoneurons in Drosophila. J Neurosci. 40:JN-RM-0925-20. doi:10.1523/jneurosci.0925-20.2020.

Bhamidipati, A., S.A. Lewis, and N.J. Cowan. 2000. Adp Ribosylation Factor-like Protein 2 (Arl2) Regulates the Interaction of Tubulin-Folding Cofactor D with Native Tubulin. J Cell Biology. 149:1087–1096. doi:10.1083/jcb.149.5.1087.

Bodaleo, F.J., and C. Gonzalez-Billault. 2016. The Presynaptic Microtubule Cytoskeleton in Physiological and Pathological Conditions: Lessons from Drosophila Fragile X Syndrome and Hereditary Spastic Paraplegias. Frontiers Mol Neurosci. 9:60. doi:10.3389/fnmol.2016.00060.

Choudhary, C., C. Kumar, F. Gnad, M.L. Nielsen, M. Rehman, T.C. Walther, J.V. Olsen, and M. Mann. 2009. Lysine Acetylation Targets Protein Complexes and Co-Regulates Major Cellular Functions. Science. 325:834–840. doi:10.1126/science.1175371.

Chu, C.-W., F. Hou, J. Zhang, L. Phu, A.V. Loktev, D.S. Kirkpatrick, P.K. Jackson, Y. Zhao, and H. Zou. 2011. A novel acetylation of β-tubulin by San modulates microtubule polymerization via down-regulating tubulin incorporation. Mol Biol Cell. 22:448–456. doi:10.1091/mbc.e10-03-0203.

Coombes, C., A. Yamamoto, M. McClellan, T.A. Reid, M. Plooster, G.W.G. Luxton, J. Alper, J. Howard, and M.K. Gardner. 2016. Mechanism of microtubule lumen entry for the α-tubulin acetyltransferase enzyme αTAT1. Proc National Acad Sci. 113:E7176–E7184. doi:10.1073/pnas.1605397113.

Coombes, C.E., H.A.J. Saunders, A.G. Mannava, D.M. Johnson-Schlitz, T.A. Reid, S. Parmar, M. McClellan, C. Yan, S.L. Rogers, J.Z. Parrish, M. Wagenbach, L. Wordeman, J. Wildonger, and M.K. Gardner. 2020. Non-enzymatic Activity of the α-Tubulin Acetyltransferase αTAT Limits Synaptic Bouton Growth in Neurons. Curr Biol. 30:610-623.e5. doi:10.1016/j.cub.2019.12.022.

Haggarty, S.J., K.M. Koeller, J.C. Wong, C.M. Grozinger, and S.L. Schreiber. 2003. Domain-selective small-molecule inhibitor of histone deacetylase 6 (HDAC6)-mediated tubulin deacetylation. Proc National Acad Sci. 100:4389–4394. doi:10.1073/pnas.0430973100.

Hansen, B.K., R. Gupta, L. Baldus, D. Lyon, T. Narita, M. Lammers, C. Choudhary, and B.T. Weinert. 2019. Analysis of human acetylation stoichiometry defines mechanistic constraints on protein regulation. Nat Commun. 10:1055. doi:10.1038/s41467-019-09024-0.

Howes, S.C., G.M. Alushin, T. Shida, M.V. Nachury, and E. Nogales. 2014. Effects of tubulin acetylation and tubulin acetyltransferase binding on microtubule structure. Mol Biol Cell. 25:257–266. doi:10.1091/mbc.e13-07-0387.

Hubbert, C., A. Guardiola, R. Shao, Y. Kawaguchi, A. Ito, A. Nixon, M. Yoshida, X.-F. Wang, and T.-P. Yao. 2002. HDAC6 is a microtubule-associated deacetylase. Nature. 417:455. doi:10.1038/417455a.

Hummel, T., K. Krukkert, J. Roos, G. Davis, and C. Klämbt. 2000. Drosophila Futsch/22C10 Is a MAP1B-like Protein Required for Dendritic and Axonal Development. Neuron. 26:357–370. doi:10.1016/s0896-6273(00)81169-1.

Janke, C., and M.M. Magiera. 2020. The tubulin code and its role in controlling microtubule properties and functions. Nat Rev Mol Cell Bio. 21:307–326. doi:10.1038/s41580-020-0214-3.

Jenkins, B.V., H.A.J. Saunders, H.L. Record, D.M. Johnson-Schlitz, and J. Wildonger. 2017. Effects of mutating α-tubulin lysine 40 on sensory dendrite development. J Cell Sci. doi:10.1242/jcs.210203.

Jin, S., L. Pan, Z. Liu, Q. Wang, Z. Xu, and Y.Q. Zhang. 2009. Drosophila Tubulin-specific chaperone E functions at neuromuscular synapses and is required for microtubule network formation. Development. 136:1571–1581. doi:10.1242/dev.029983.

Kalebic, N., S. Sorrentino, E. Perlas, G. Bolasco, C. Martinez, and P.A. Heppenstall. 2013. αTAT1 is the major α-tubulin acetyltransferase in mice. Nat Commun. 4. doi:10.1038/ncomms2962.

Kaluza, D., J. Kroll, S. Gesierich, T. Yao, R.A. Boon, E. Hergenreider, M. Tjwa, L. Rössig, E. Seto, H.G. Augustin, A.M. Zeiher, S. Dimmeler, and C. Urbich. 2011. Class IIb HDAC6 regulates endothelial cell migration and angiogenesis by deacetylation of cortactin. Embo J. 30:4142–4156. doi:10.1038/emboj.2011.298.

Kamiyama, D., S. Sekine, B. Barsi-Rhyne, J. Hu, B. Chen, L.A. Gilbert, H. Ishikawa, M.D. Leonetti, W.F. Marshall, J.S. Weissman, and B. Huang. 2016. Versatile protein tagging in cells with split fluorescent protein. Nat Commun. 7:11046. doi:10.1038/ncomms11046.

Kelliher, M.T., Y. Yue, A. Ng, D. Kamiyama, B. Huang, K.J. Verhey, and J. Wildonger. 2018. Autoinhibition of kinesin-1 is essential to the dendrite-specific localization of Golgi outposts. J Cell Biol. jcb.201708096. doi:10.1083/jcb.201708096.

Kim, G.-W., L. Li, M. Gorbani, L. You, and X.-J. Yang. 2013. Mice Lacking α-Tubulin Acetyltransferase 1 Are Viable but Display α-Tubulin Acetylation Deficiency and Dentate Gyrus Distortion. J Biol Chem. 288:20334–20350. doi:10.1074/jbc.m113.464792.

Knossow, M., V. Campanacci, L.A. Khodja, and B. Gigant. 2020. The mechanism of tubulin assembly into microtubules: insights from structural studies. Iscience. 23:101511. doi:10.1016/j.isci.2020.101511.

Lepicard, S., B. Franco, F. de Bock, and M.-L. Parmentier. 2014. A Presynaptic Role of Microtubule-Associated Protein 1/Futsch in Drosophila: Regulation of Active Zone Number and Neurotransmitter Release. J Neurosci. 34:6759–6771. doi:10.1523/jneurosci.4282-13.2014.

L’Hernault, S.W., and J.L. Rosenbaum. 1985. Chlamydomonas .alpha.-tubulin is posttranslationally modified by acetylation on the .epsilon.-amino group of a lysine. Biochemistry-us. 24:473–478. doi:10.1021/bi00323a034.

Liu, N., Y. Xiong, S. Li, Y. Ren, Q. He, S. Gao, J. Zhou, and W. Shui. 2015a. New HDAC6-mediated deacetylation sites of tubulin in the mouse brain identified by quantitative mass spectrometry. Sci Reports. 5:16869. doi:10.1038/srep16869.

Liu, N., Y. Xiong, Y. Ren, L. Zhang, X. He, X. Wang, M. Liu, D. Li, W. Shui, and J. Zhou. 2015b. Proteomic Profiling and Functional Characterization of Multiple Post-Translational Modifications of Tubulin. J Proteome Res. 14:3292–3304. doi:10.1021/acs.jproteome.5b00308.

Lundby, A., K. Lage, B.T. Weinert, D.B. Bekker-Jensen, A. Secher, T. Skovgaard, C.D. Kelstrup, A. Dmytriyev, C. Choudhary, C. Lundby, and J.V. Olsen. 2012. Proteomic Analysis of Lysine Acetylation Sites in Rat Tissues Reveals Organ Specificity and Subcellular Patterns. Cell Reports. 2:419–431. doi:10.1016/j.celrep.2012.07.006.

Ma, X.-X., X. Li, P. Yi, C. Wang, J. Weng, L. Zhang, X. Xu, H. Sun, S. Feng, K. Liu, R. Chen, S. Du, X. Mao, X. Zeng, L.-Y. Zhang, M. Liu, B.-S. Tang, X. Zhu, S. Jin, and J.-Y. Liu. 2017. PiT2 regulates neuronal outgrowth through interaction with microtubule-associated protein 1B. Sci Reports. 7:17850. doi:10.1038/s41598-017-17953-3.

Mao, C.-X., X. Wen, S. Jin, and Y.Q. Zhang. 2017. Increased acetylation of microtubules rescues human tau-induced microtubule defects and neuromuscular junction abnormalities in Drosophila. Dis Model Mech. 10:dmm.028316. doi:10.1242/dmm.028316.

McLaughlin, C.N., I.V. Nechipurenko, N. Liu, and H.T. Broihier. 2016. A Toll receptor–FoxO pathway represses Pavarotti/MKLP1 to promote microtubule dynamics in motoneurons. J Cell Biology. 214:459–474. doi:10.1083/jcb.201601014.

Miskiewicz, K., L.E. Jose, W.M. Yeshaw, J.S. Valadas, J. Swerts, S. Munck, F. Feiguin, B. Dermaut, and P. Verstreken. 2014. HDAC6 Is a Bruchpilot Deacetylase that Facilitates Neurotransmitter Release. Cell Reports. 8:94–102. doi:10.1016/j.celrep.2014.05.051.

Nechipurenko, I.V., and H.T. Broihier. 2012. FoxO limits microtubule stability and is itself negatively regulated by microtubule disruption. J Cell Biology. 196:345–362. doi:10.1083/jcb.201105154.

Neumann, B., and M.A. Hilliard. 2014. Loss of MEC-17 Leads to Microtubule Instability and Axonal Degeneration. Cell Reports. 6:93–103. doi:10.1016/j.celrep.2013.12.004.

Nithianantham, S., S. Le, E. Seto, W. Jia, J. Leary, K.D. Corbett, J.K. Moore, and J. Al-Bassam. 2015. Tubulin cofactors and Arl2 are cage-like chaperones that regulate the soluble αβ-tubulin pool for microtubule dynamics. Elife. 4. doi:10.7554/elife.08811.

Okumura, M., C. Sakuma, M. Miura, and T. Chihara. 2015. Linking Cell Surface Receptors to Microtubules: Tubulin Folding Cofactor D Mediates Dscam Functions during Neuronal Morphogenesis. J Neurosci. 35:1979–1990. doi:10.1523/jneurosci.0973-14.2015.

Osseni, A., A. Ravel-Chapuis, J.-L. Thomas, V. Gache, L. Schaeffer, and B.J. Jasmin. 2020. HDAC6 regulates microtubule stability and clustering of AChRs at neuromuscular junctions. J Cell Biol. 219. doi:10.1083/jcb.201901099.

Penazzi, L., L. Bakota, and R. Brandt. 2016. International Review of Cell and Molecular Biology. undefined. 321:89–169. doi:10.1016/bs.ircmb.2015.09.004.

Portran, D., L. Schaedel, Z. Xu, M. Théry, and M.V. Nachury. 2017. Tubulin acetylation protects long-lived microtubules against mechanical ageing. Nat Cell Biol. 19:391–398. doi:10.1038/ncb3481.

Raff, E.C. 1984. Genetics of microtubule systems. J Cell Biology. 99:1–10. doi:10.1083/jcb.99.1.1.

Roos, J., T. Hummel, N. Ng, C. Klämbt, and G.W. Davis. 2000. Drosophila Futsch Regulates Synaptic Microtubule Organization and Is Necessary for Synaptic Growth. Neuron. 26:371–382. doi:10.1016/s0896-6273(00)81170-8.

Serna, M., G. Carranza, J. Martín-Benito, R. Janowski, A. Canals, M. Coll, J.C. Zabala, and J.M. Valpuesta. 2015. The structure of the complex between α-tubulin, TBCE and TBCB reveals a tubulin dimer dissociation mechanism. J Cell Sci. 128:1824–1834. doi:10.1242/jcs.167387.

Shi, Q., Y.Q. Lin, A. Saliba, J. Xie, G.G. Neely, and S. Banerjee. 2019. Tubulin Polymerization Promoting Protein, Ringmaker, and MAP1B Homolog Futsch Coordinate Microtubule Organization and Synaptic Growth. Front Cell Neurosci. 13:192. doi:10.3389/fncel.2019.00192.

Shida, T., J.G. Cueva, Z. Xu, M.B. Goodman, and M.V. Nachury. 2010. The major α-tubulin K40 acetyltransferase αTAT1 promotes rapid ciliogenesis and efficient mechanosensation. Proc Natl Acad Sci. 107:21517–21522. doi:10.1073/pnas.1013728107.

Szasz, J., M.B. Yaffe, M. Elzinga, G.S. Blank, and H. Sternlicht. 1986. Microtubule assembly is dependent on a cluster of basic residues in .alpha.-tubulin. Biochemistry-us. 25:4572–4582. doi:10.1021/bi00364a018.

Szasz, J., M.B. Yaffe, and H. Sternlicht. 1993. Site-directed mutagenesis of alpha-tubulin. Reductive methylation studies of the Lys 394 region. Biophys J. 64:792–802. doi:10.1016/s0006-3495(93)81440-1.

Szyk, A., A.M. Deaconescu, J. Spector, B. Goodman, M.L. Valenstein, N.E. Ziolkowska, V. Kormendi, N. Grigorieff, and A. Roll-Mecak. 2014. Molecular Basis for Age-Dependent Microtubule Acetylation by Tubulin Acetyltransferase. Cell. 157:1405–1415. doi:10.1016/j.cell.2014.03.061.

Topalidou, I., C. Keller, N. Kalebic, K.C.Q. Nguyen, H. Somhegyi, K.A. Politi, P. Heppenstall, D.H. Hall, and M. Chalfie. 2012. Genetically Separable Functions of the MEC-17 Tubulin Acetyltransferase Affect Microtubule Organization. Curr Biol. 22:1057–1065. doi:10.1016/j.cub.2012.03.066.

Valenzuela-Fernández, A., J.R. Cabrero, J.M. Serrador, and F. Sánchez-Madrid. 2008. HDAC6: a key regulator of cytoskeleton, cell migration and cell–cell interactions. Trends Cell Biol. 18:291–297. doi:10.1016/j.tcb.2008.04.003.

Wang, Y., M. Lobb-Rabe, J. Ashley, V. Anand, and R.A. Carrillo. 2021. Structural and functional synaptic plasticity induced by convergent synapse loss in the Drosophila neuromuscular circuit. J Neurosci. 41:JN-RM-1492-20. doi:10.1523/jneurosci.1492-20.2020.

Wei, D., N. Gao, L. Li, J.-X. Zhu, L. Diao, J. Huang, Q.-J. Han, S. Wang, H. Xue, Q. Wang, Q.-F. Wu, X. Zhang, and L. Bao. 2017. α-Tubulin Acetylation Restricts Axon Overbranching by Dampening Microtubule Plus-End Dynamics in Neurons. Cereb Cortex New York N Y 1991. 1–15. doi:10.1093/cercor/bhx225.

Weinert, B.T., S.A. Wagner, H. Horn, P. Henriksen, W.R. Liu, J.V. Olsen, L.J. Jensen, and C. Choudhary. 2011. Proteome-wide mapping of the Drosophila acetylome demonstrates a high degree of conservation of lysine acetylation. Sci Signal. 4:ra48. doi:10.1126/scisignal.2001902.

Xiong, Y., K. Zhao, J. Wu, Z. Xu, S. Jin, and Y.Q. Zhang. 2013. HDAC6 mutations rescue human tau-induced microtubule defects in Drosophila. Proc National Acad Sci. 110:4604–4609. doi:10.1073/pnas.1207586110.

Xu, Z., L. Schaedel, D. Portran, A. Aguilar, J. Gaillard, M.P. Marinkovich, M. Théry, and M.V. Nachury. 2017. Microtubules acquire resistance from mechanical breakage through intralumenal acetylation. Science. 356:328–332. doi:10.1126/science.aai8764.

Yan, C., F. Wang, Y. Peng, C.R. Williams, B. Jenkins, J. Wildonger, H.-J. Kim, J.B. Perr, J.C. Vaughan, M.E. Kern, M.R. Falvo, E.T. O’Brien, R. Superfine, J.C. Tuthill, Y. Xiang, S.L. Rogers, and J.Z. Parrish. 2018. Microtubule Acetylation Is Required for Mechanosensation in Drosophila. Cell Reports. 25:1051-1065.e6. doi:10.1016/j.celrep.2018.09.075.

Yin, S., C. Zeng, M. Hari, and F. Cabral. 2013. Paclitaxel resistance by random mutagenesis of α-tubulin. Cytoskeleton. 70:849–862. doi:10.1002/cm.21154.

Zou, B., H. Yan, F. Kawasaki, and R.W. Ordway. 2008. MAP1 structural organization in Drosophila: in vivo analysis of FUTSCH reveals heavy-and light-chain subunits generated by proteolytic processing at a conserved cleavage site. Biochem J. 414:63–71. doi:10.1042/bj20071449.

